# The Impact of a Dynamic Working Environment on Human Gut Microbiota

**DOI:** 10.1101/2021.04.19.440557

**Authors:** Lu Ling, Jun Zhou, Qianlong Meng, Ziran Zhang, Wenkun Li, Zilu Cui, Kuiliang Liu, Fanxin Zeng, Jing Wu, Jing Wang

## Abstract

Gut microbiota dysbiosis is associated with a variety of diseases, such as inflammatory bowel disease (IBD) and irritable bowel syndrome (IBS), metabolic diseases, allergic diseases, neurodevelopmental disorders and cancer. The human gut microbiota can be influenced by a variety of factors, including geography, dietary habits, living environment, age and altered lifestyle etc. This study was conducted to explore the gut microbiota compositions in officials who are in a stable working environment and train drivers who are in a dynamic working environment. Microbiota communities in the feces of 80 officials and 88 train drivers were analyzed using Illumina MiSeq sequencing targeting the V3-V4 region of 16S ribosomal RNA (rRNA) gene and ITS1 region of fungi. There were significant differences between the two groups in diversity and richness of gut microbiota, while the microbial community compositions of the two groups were similar. The relationship between gut microbiota and clinical characteristics was investigated. We found that more bacteria and fungi were positively correlated with clinical characteristics. Functional prediction analysis of the gut microbiota between the two groups by PICRUSt2 revealed significant differences between the official group and the train driver group. Elucidating these differences of the microbiome between the two groups will provide a foundation understanding of the impact of a dynamic environment on gut microbiota.

## 1. Introduction

Humans and other mammals have complex gut microbiota including archaea, bacteria, fungi, viruses, and protists [1]. This symbiosis phenomenon is developed through the exposure to a range of environmental microorganism, beginning with the birth through the vagina with specific microbiome [2]. The microbiome is usually assumed to be the bacterial microbiota and the gut microbiota is the most studied so far [3]. It has been the major topic of interest, due to the large amounts (∼10^13^-10^14^) and great importance of bacteria found in our gastrointestinal tract (GI) [4]. The gastrointestinal tract is mainly composed of strict anaerobe, the number of which is two to three times more than that of facultative anaerobe and aerobic bacteria [5]. Commensal fungi that normally inhabit our bodies have been much less studied, as fungi form the minority of the total commensal organisms in humans [6].

The human gut microbiota participates in multiple host activities including nutrient absorption, metabolic and immune regulations. However, the role of many fungi remains unclear. The alterations in the composition of microbiota can have adverse effects on host health, especially the dysbiosis. A growing body of clinical evidence suggests that gut microbiota dysbiosis is associated with a variety of immune, metabolic, and neurological diseases in the intestinal and extraintestinal tracts. Pathological states associated with altered microbiota include inflammatory bowel disease (IBD) and irritable bowel syndrome (IBS), as well as metabolic diseases such as obesity and diabetes mellitus, allergic diseases, neurodevelopmental disorders and cancer [7]. In addition, many pathogenic fungi are symbiotic organisms in our bodies that normally do no harm but have the potential to cause disease. For example, *candida albicans*, which cause systemic candidiasis in immunocompromised patients, are a normal part of the intestinal flora [8]. What causes these normally non-pathogenic fungi to cause disease is still a hot topic of debate [4]. The definition of mycobiome broadens the scope of the study of the human microbiome.

Microbiota can cause a variety of diseases through different mechanisms, including production of harmful decomposition metabolites, overgrowth, persistent inflammation, or the ability to provide support for pathogens [9]. The human gut microbiota can be influenced by a variety of factors, including geography, dietary habits, living environment, age and altered lifestyle etc. In conclusion, it is important to understand the composition, function and impact factors of gut microbiota for maintaining human health and monitoring the occurrence and development of diseases. The officials and train drivers are two groups of people working in different environment. The train drivers work in a changing environment compared with the officials. Elucidating these differences of the microbiome between the two groups will provide a foundation understanding of the impact of environment on gut microbiota.

## 2. Material and methods

### 2.1. Sampling and information collection

A total of 168 employees in China Railway Corporation were recruited in this study, including 80 officials and 88 train drivers as two groups of subjects. Exclusion criterions for subjects were serious chronic illness (e.g., diabetes, heart failure, cancer or autoimmune diseases). All enrolled subjects provided written informed consent. Our research received the approval from the ethics committee, Shijitan Hospital, Capital Medical University, Beijing, China. The mean age of the two groups was 48.65 (±5.28) years and 44.99 (±3.67) years, respectively. The majority of railway workers are male, and the proportion of male in each group is 83.1% and 100%, respectively. A fecal sample was collected from each subject. Fecal samples were frozen immediately after sampling and stored at -80°C.

All the subjects were surveyed by questionnaires, which included general information such as age, gender, body mass index (BMI), marriage, lifestyles, dietary habits and health status. All subjects’ weight and height were measured, and BMI was calculated from weight in kilograms divided by height in meters squared. The frequency of physical exercise per week was defined as ⩽1 day, 2-3 days and ⩾ 4 days respectively. The labor intensity was classified as light-degree, moderate-degree, heavy-degree and extremely heavy-degree. The frequency of diet was defined as none, occasional, frequent and daily respectively. A standard question was completed by the same physician through a face-to-face interview.

### 2.2. DNA extraction, 16S rRNA gene and ITS amplicon and sequencing

Total fecal genomic DNA was extracted using QIAamp DNA Stool Mini Kit (Qiagen, Hilden, Germany) according to the manufacturer’s instruction. The integrity and size of DNA was assessed by 1% agarose gel electrophoresis. The bacterial 16S rRNA gene fragments (V3-V4) were amplified using universal primers 341F (5’-ACTCCTACGGGAGGCAGCAG-3’) and 806R (5’-GGACTACHVGGGTWTCTAAT-3’), and the fungal internal transcribed spacer regions 1 (ITS1) was amplified using primers ITS1F (CTTGGTCATTTAGAGGAAGTAA) and ITS2R (GCTGCGTTCTTCATCGATGC) with the following PCR conditions: 30 s at 95 °C, 30 s at 55 °C, and 45 s at 72 °C for 27 cycles.The amplicons were sequenced on an Illumina MiSeq platform.

After demultiplexing, the sequences were merged with FLASH (v1.2.11) and quality filtered with fastp (version 0.19.6). Then the high-quality sequences were de-noised using DADA2 plugin in the Qiime2 (version 2020.2) pipeline with recommended parameters, which obtains single nucleotide resolution based on error profiles within samples. DADA2 denoised sequences are usually defined amplicon sequence variants (ASVs). To analyze the sequencing depth on alpha and beta diversity, each sample sequences was rarefied to same level according to the minimum sequence number of samples. Taxonomic assignment of ASVs was performed by the Naive bayes consensus taxonomy classifier implemented in Qiime2. The databases of 16S and ITS are SILVA 16S rRNA database (v138) and UNITE 8.0 respectively. The functional prediction of the community was carried out by the phylogenetic investigation of communities by reconstruction of unobserved states 2 (PICRUSt2).

### 2.3. Statistical analysis

SPSS 25.0, GraphPad Prism 8.0, Cytoscape 3.7.2, STAMP and R version 3.5.1 software were used for statistical analysis and plotting. Normal distribution of continuous variables were reported as mean ± standard deviation, and analyzed by Student’s *t* test, while non-normal distribution of continuous variables were reported as median with interquartile ranges and analyzed by Mann-Whitney *U* test. Categorical variables were reported as percentages and analyzed by Pearson’s Chi-square test. The difference of gut microbiota between the two groups was compared by Wilcoxon rank-sum test. The correlation between clinical characteristics and intestinal microbiota was analyzed by Spearman’s rank correlation. *P* < 0.05 was considered statistically significant.

## 3. Results

### 3.1 The basic statistics and clinical characteristics of the subjects

A total of 168 railway workers, including 88 officials and 80 train drivers, were included in bacterial analysis. The average age of the two groups was 48.60 ± 5.31 years and 44.96 ± 3.72 years, respectively. Since the majority of railway workers are male, and the proportion of male in each group is 84.09% and 100%, respectively. The average BMI of the two groups was 27.04 ± 4.69 kg/m^2^ and 26.00 ± 3.32 kg/m^2^, respectively. There were significant differences between the two groups in age, gender, sitting, night watch, night watch frequency, labor intensity and going out with the train (*P* < 0.05). The rest information was shown in Table 1.

**Table 1.**
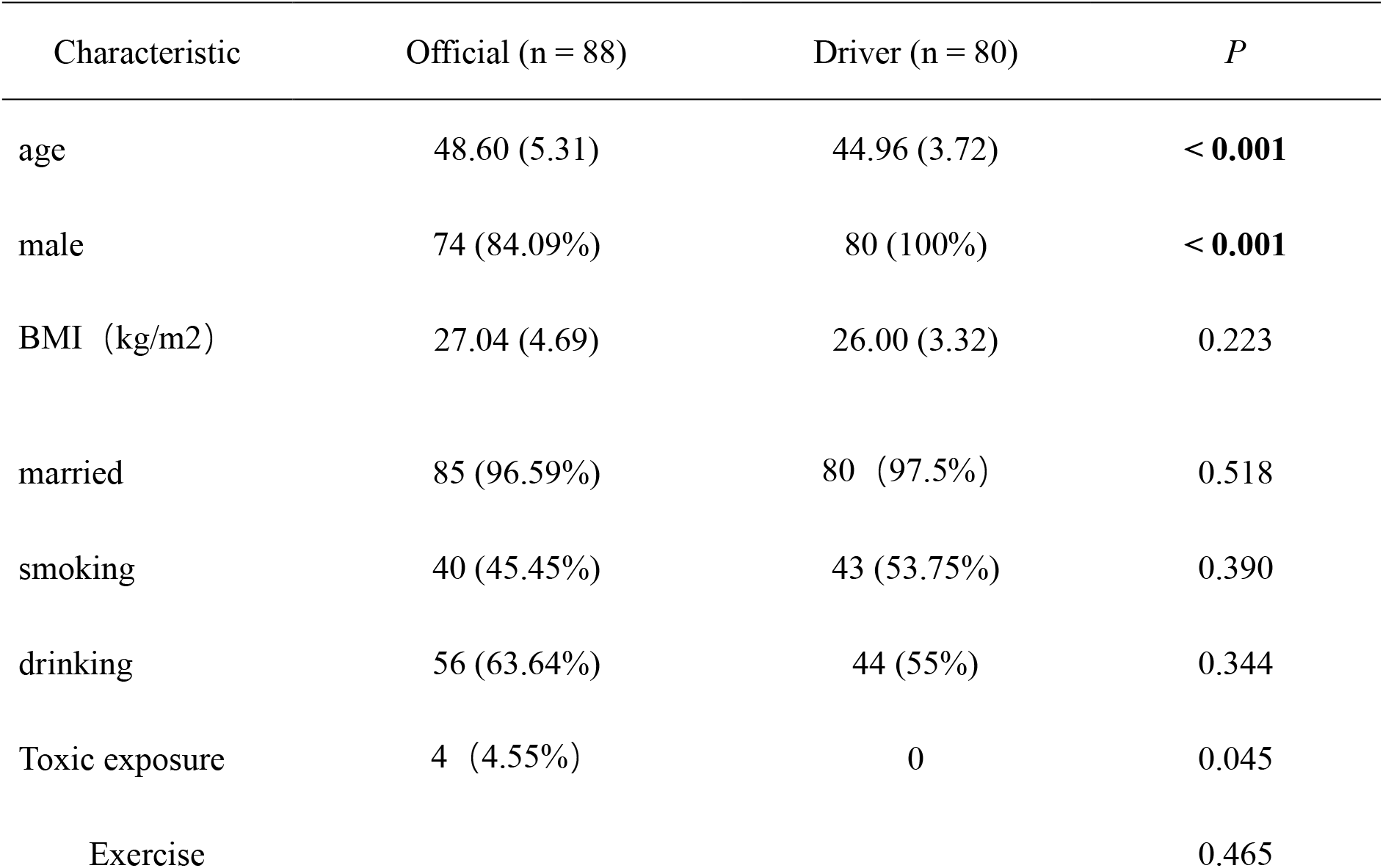
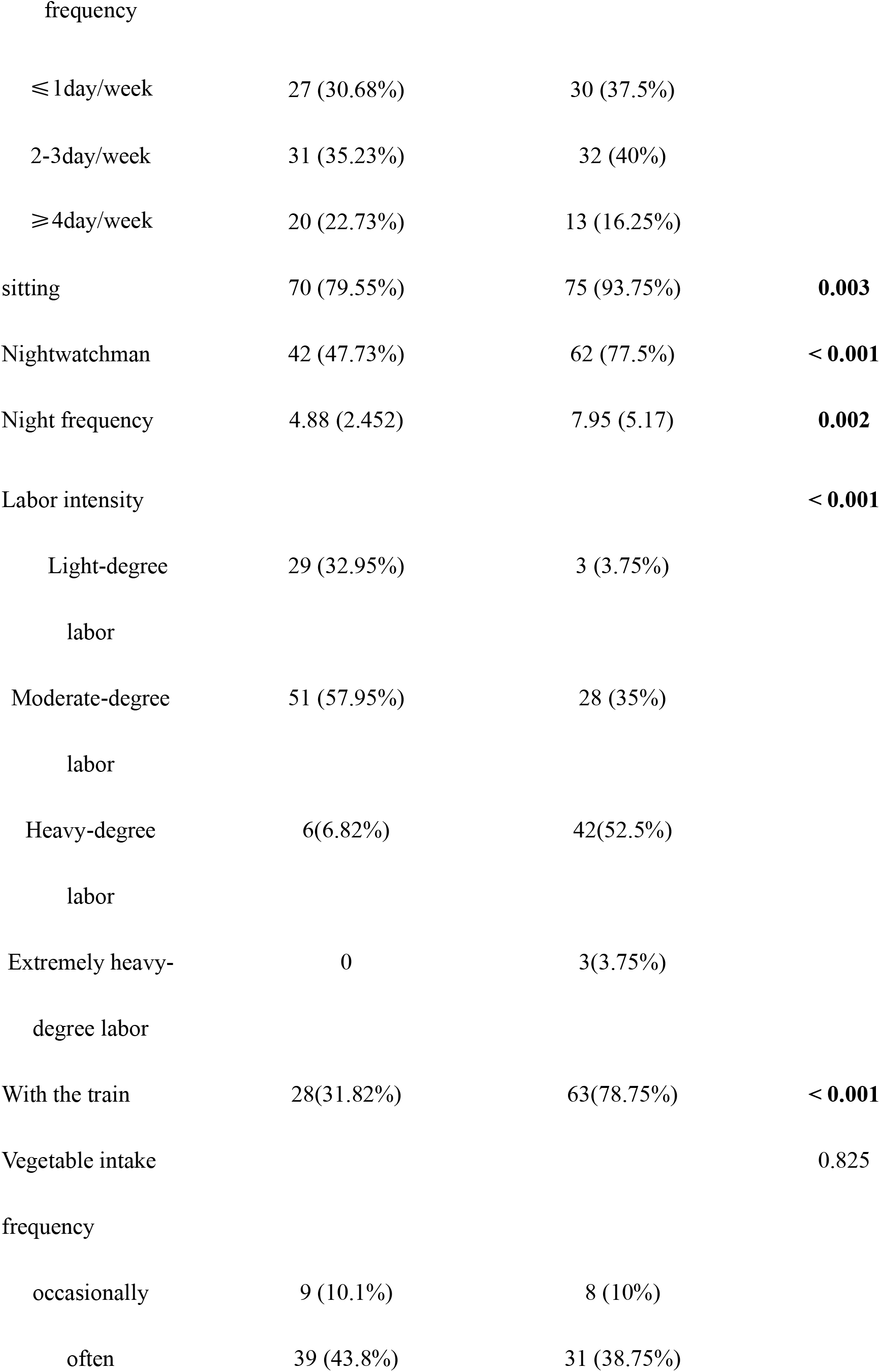
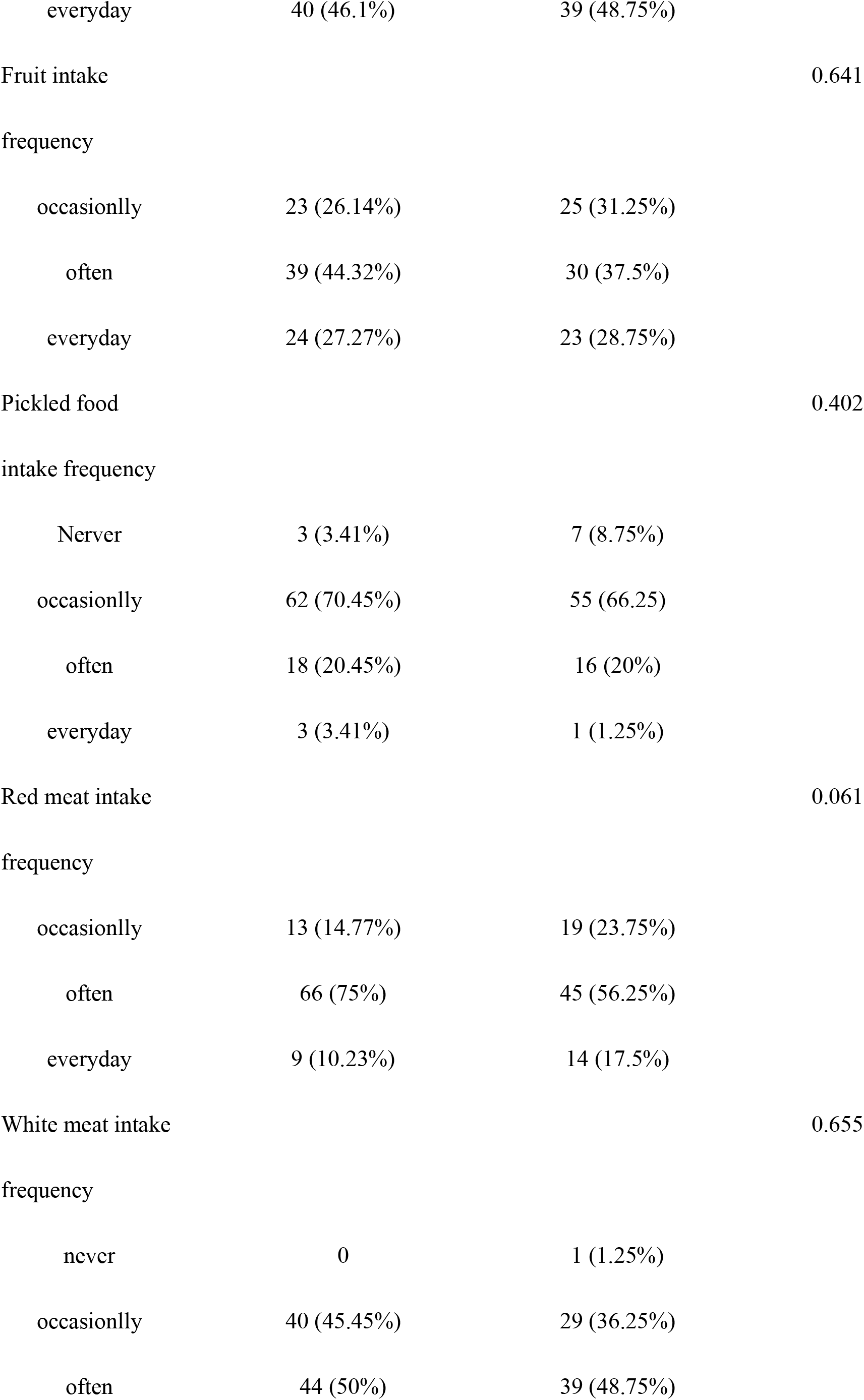
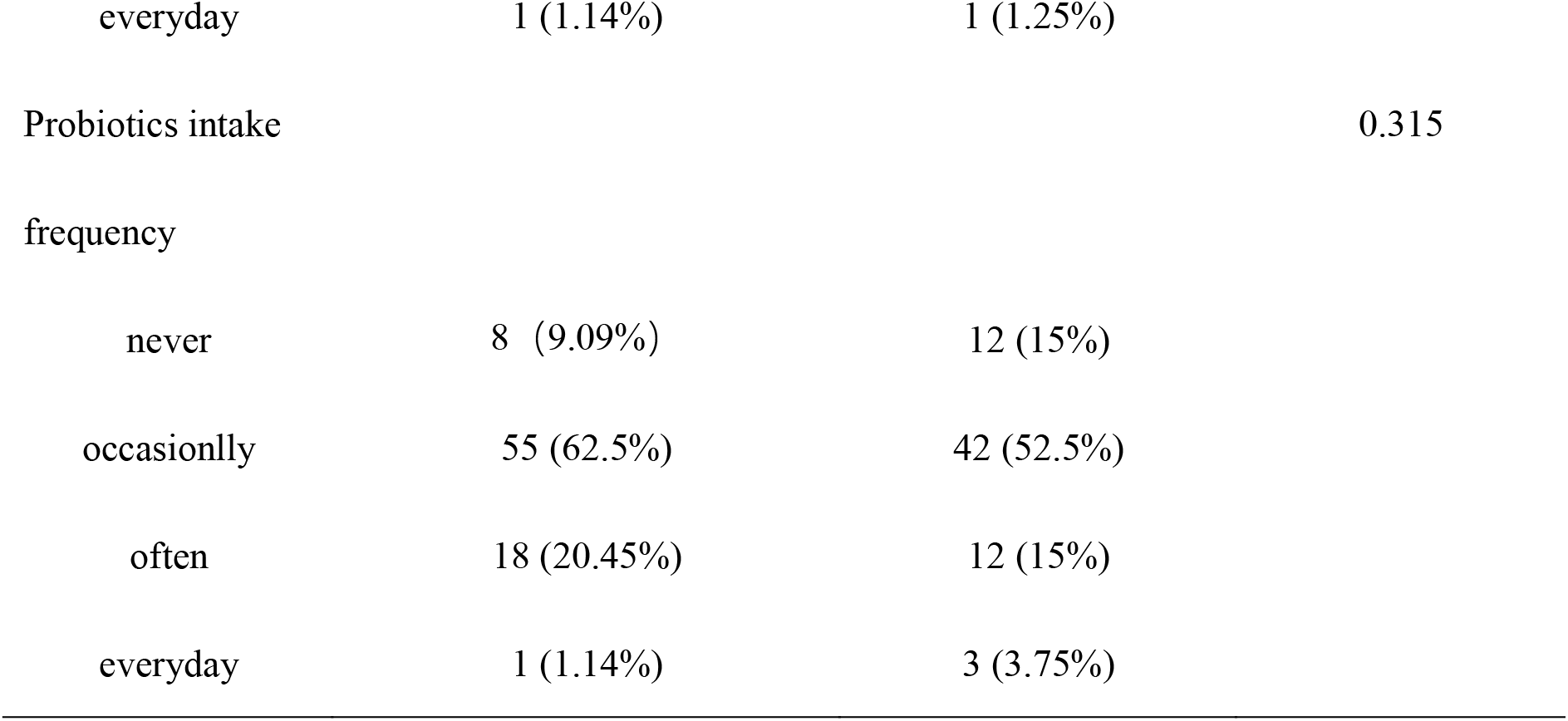
Baseline statistics and clinical characteristics of the subjects

### 3.2 Diversity analysis for gut microbiota between officials and train drivers

A total of 32,346,775 valid sequences were obtained by 16S rRNA gene sequencing, and 20,434,052 sequences were obtained by ITS sequencing. The rank-abundance curves for bacterial and fungal gut microbiota showed that species uniform was similar between officials and train drivers (Fig.S1 A, B). Besides, the numbers of total genus by Pan analyses for 16S and ITS were similar between the two groups (Fig.S1 C, D). There was no difference in the coverage rates between the two groups (Figure 1). The coverage rates of both groups were about 1, indicating that the sequencing depth has covered all the species in the sample. The alpha diversity indexes including Ace, Chao, Shannon, Sobs in the officials were significantly higher than those of the train drivers, while there was no significantly difference in Simpson index. These indicated that the richness and diversity of the gut microbiota in officials were significantly higher than those of train drivers. Nonmetric Multidimensional Scaling (NMDS) analysis showed no significant difference in bacterial and fungal community compositions between the two groups (Figure1 C,D).

**Fig. 1.**
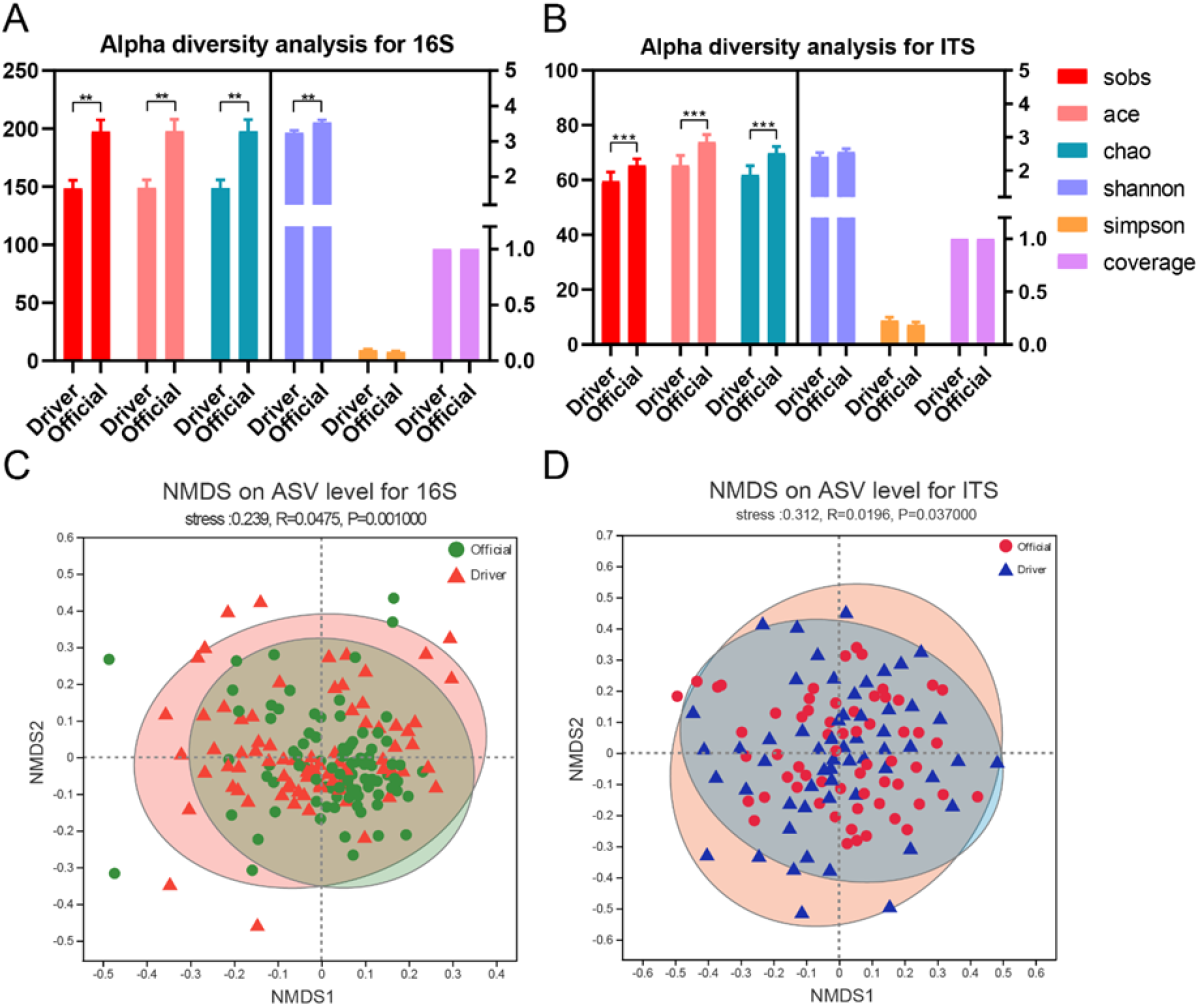
Diversity analysis for gut microbiota. A, Alpha diversity analysis for the bacterial gut microbiota; B, Alpha diversity analysis for the fungal gut microbiota; C, NMDS analysis on ASV level for the bacterial gut microbiota; D, NMDS analysis on ASV level for the fungal gut microbiota. The higher the similarity between samples, the more concentrated they are in the figure. * *P* < 0.05, ** *P* < 0.01.

### 3.3. Microbiota composition of official and train driver groups

In 16S rRNA gene sequencing, the Venn plot based on the level of ASV showed that there were 1169 ASVs in common in the official group and the train driver group, 5625 special ASVs in the official group and 2909 special ASVs in the train driver group (Figure 2A); While in ITS sequencing, there were 445 ASVs in common in the official group and the train driver group, 1764 special ASVs in the official group and 1511 special ASVs in the train driver group (Figure 2B).

**Fig. 2.**
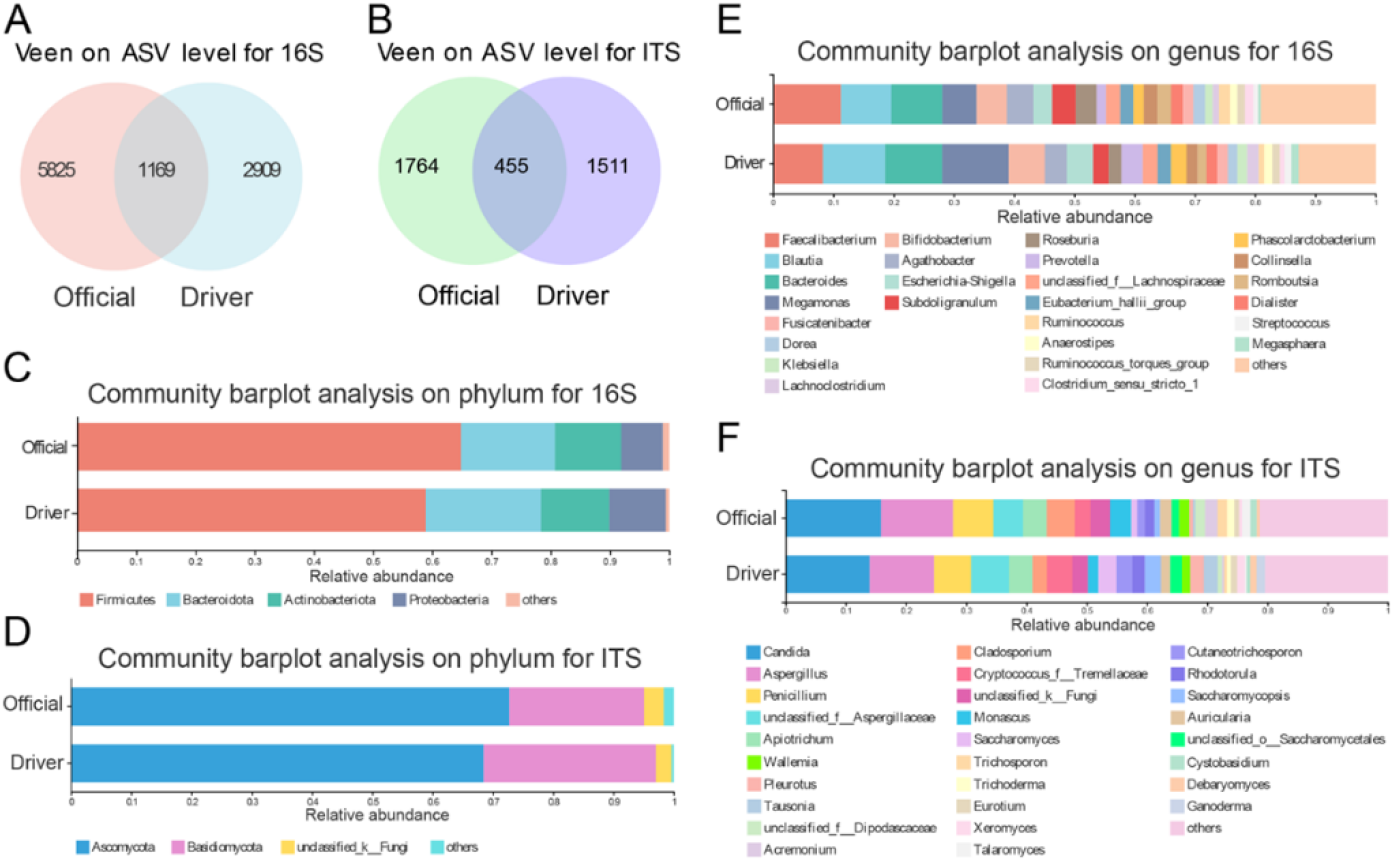
Microbiological composition for the gut microbiota. A, Venn analysis on ASV level for the bacterial gut microbiota; B, Venn analysis on ASV level for the fungal gut microbiota; Different groups in the figure are represented by different colors, and the numbers in the figure represent specific or common ASV numbers. The overlapping region represents the number of ASVs common to different groups, while the non-overlapping region represents the number of ASVs unique to different groups. C, Community barplot analysis on phylum level for the bacterial gut microbiota; D, Community barplot analysis on phylum level the fungal gut microbiota; E, Community barplot analysis on genus level for the bacterial gut microbiota; F, Community barplot analysis on genus level for the fungal gut microbiota.

The 16S rRNA gene sequencing analysis showed that, the fecal bacteria of official group and train driver group were both composed of Fimicutes, Bacteroidota, Actinobateriota, and Protecbacteria, and Fimicutes accounted for more than 70% of the total fecal microflora on phylum level (Figure 2C). The relative abundance of Fimicutes in fecal bacteria of the official group was significantly higher than that of the train driver group, while the relative abundance of Bacteroidota and Protecbacteria in fecal microbiota of the train driver group was significantly higher than that of the officials group. On genus level, the fecal bacteria of the official group and the train driver group were both composed of *Faecalibacterium, Blautia, Bacteroides, Megamonas, Bifidobacterium, Agathobacter, Escherichia*-*Shigella, Subdoligranulum, Roseburia, Preverella Votella, Lachnospiracee, Eubacterium Hali Group*, etc (Figure 2E).

The ITS sequencing analysis showed that, the fecal fungi of the official group and the train driver group were both mainly composed of *Ascomycota* and *Basidiomycota* on phylum level (Figure 2D). The relative abundance of *Ascomycota* in the official group was significantly higher than that in the train driver group, while the relative abundance of *Basidiomycota* in the train driver group was significantly higher than that in the official group. On genus level, the two groups were both composed of *Candida, Aspergillus, Penicillium, Apiotrichum, Cladosporium, Cryptococcus, Monascus*, etc (Figure 2F). The relative abundance of *Candida* in the official group was significantly higher than that in the train driver group.

### 3.4 Microbial difference analysis between official group and train driver group

By using Linear discriminant analysis (LDA) effect size (LEfSe) analysis to compare the fecal microbiota between the official and train driver groups, we used a LDA score cutoff of 3.0 to determine important taxonomic differences between the official and train driver groups. The results showed that there were significant differences in fecal microbiota between the official group and the train driver group based on LefSe analysis (Figure 3A, B, C, D). For the bacterial gut microbiota, the relative abundances of *Clostridia, Oscillospirales, Ruminococcaceae*, Firmicutes, *Faecalibacterium, Subdoligranulum, Roseburia, Ruminococcus, Oscillospiraceae, Clostridia*_*UCG*_014, and *christensenellales* etc. in the official group were significantly higher than those of the train driver group (*P* < 0.05), while the relative abundances of *Negativicutes, Megamonas, Ruminococcus*_*gnavus_group and Lachnoclostridium* in the train driver group were significantly higher than that of the official group (Figure 3A). For the fungal gut microbiota, the relative abundance of *Aspergillus, Monascus, Cladosporium, Dipodascaceae, Symmetrospora, Cystobasidiomycetes*_*ord*_*Incertae*_*sedis, Symmetrosporaceae, Kazachstania*, etc. in the official group was significantly higher than that in the train driver group.The relative abundance of *Cutaneotrichosporon, Cutaneotrichosporon*_*cutaneum, Aspergillus*_*halophilicus, Cryptococcus*_*uniguttulatus, Coniochaetaceae, Pestalotiopsis*_*camelliae, Coniochaetaceae, Peatalotiopsis, Corallomycetella, Corallomycetella*_*elegans, Zygosaccharomyces*_*bisporus, Trichmonascus*_*sp* in the train driver group was significantly higher than that in the official group (Figure 3C). The abundance of specific different species in LEfSe analysis were showed in Figure S2.

**Fig. 3.**
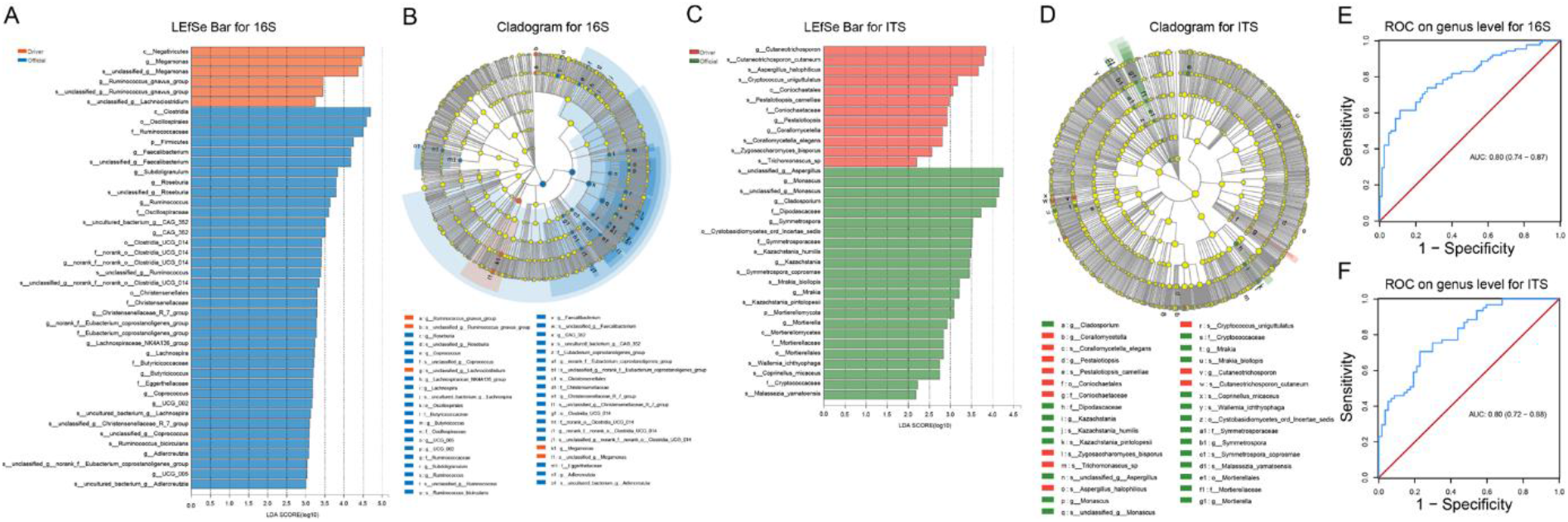
Analysis of Microbiota differences for gut microbiota between the two groups. A, LDA analysis results for the bacterial gut microbiota; B, The cladogram analyzed LEfSe of the bacterial gut microbiota; C, LDA analysis results for the fungal gut microbiota; D, The cladogram analyzed LEfSe of the fungal gut microbiota; E, Difference in bacterial gut microbiota were used to discriminate the two groups; F, Difference in fungal gut microbiota were used to discriminate the two groups.

At genus level, the bacteria and fungi screened out from the LEfSe analysis were used to distinguish between official group and train driver group, and the area under the curve (AUC) was 0.8 (95% confidence interval (CI): 0.74-0.87), 0.8 (95% confidence interval (CI): 0.72-0.88), respectively (Figure 3E, F).

### 3.5 Associations between fecal microbiota and clinical characteristics, and interactions between microbiota

We determined the correlation between the fecal microbiota and the clinical characteristics. More bacteria and fungi were positively correlated with clinical characteristics (Figure 4A, B).Network interactions between differential species from LEfSe analysis and other species were constructed. Network interactions can vividly show the abundance of genus between groups. Through the number of line connections, we found the genus that are more related to other members of the microflora, and then explore the biological significance of the correlation between these genera (Figure 4C, D).

**Fig. 4.**
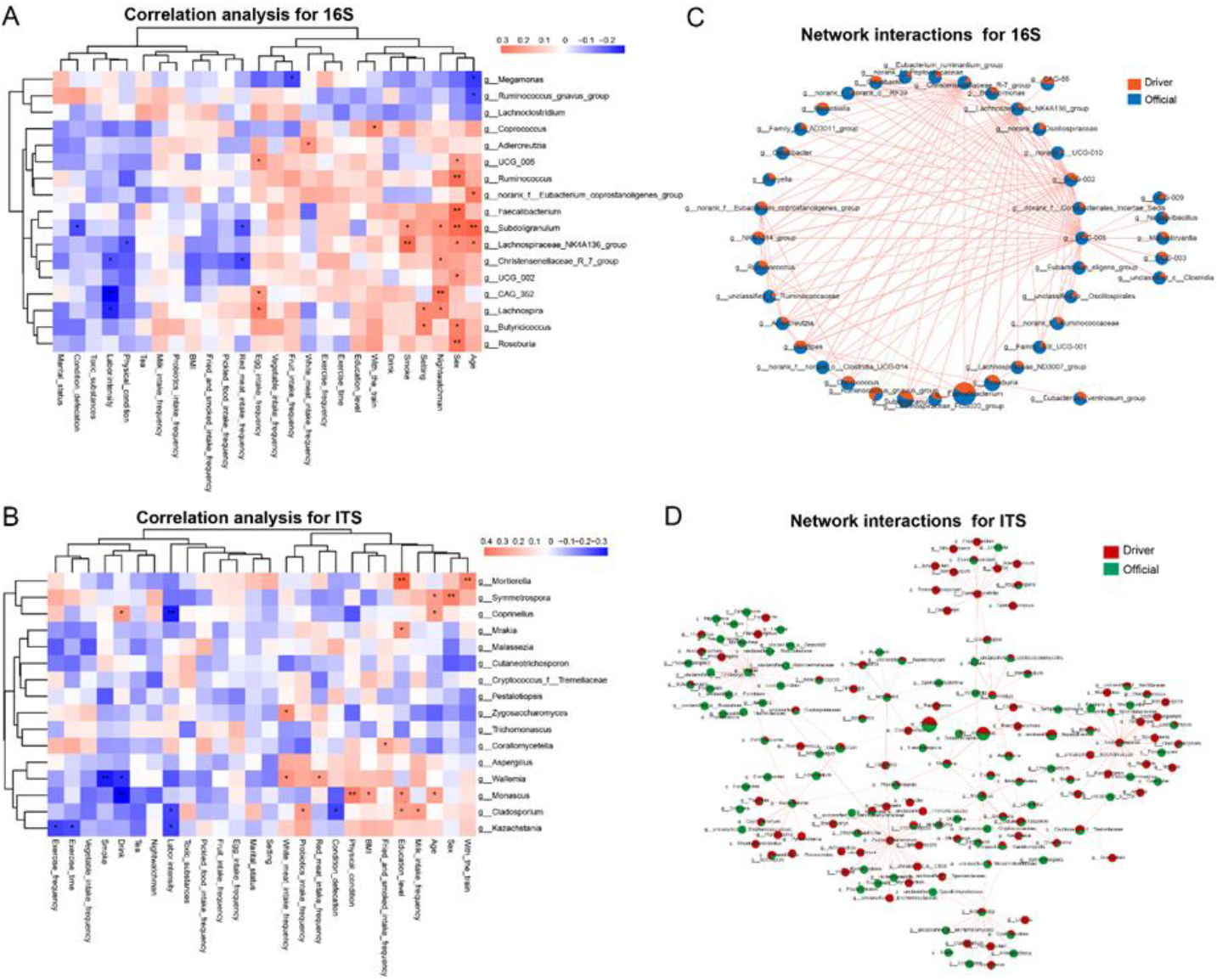
Correlation analysis for the gut microbiota. A, Correlation analysis between clinical characteristics and bacterial gut microbiota; B, Correlation analysis between clinical characteristics and fungal gut microbiota; C, Network interactions between the bacterial gut microbiota; D, Network interactions between the fungal gut microbiota. Correlation analysis was assessed by Spearman’s. * *P* < 0.05, ** *P* < 0.01. The orange line means positive regulation, green line means negative regulation. The width of the line represents the correlation coefficient between species. The size of the pie indicates the relative abundance of the genus.

### 3.7 Function prediction

In the prediction of bacterial function, there were significant differences between the official group and the train driver group in microRNAs in cancer, mismatch repair, folate biosynthesis, fatty acid metabolism, peptidoglycan biosynthesis, pentose and glucuronate interconversions (*P* ⩽0.05). The functions of folate biosynthesis and pentose and glucuronate interconversions were decreased in the official group compared with the train driver group. The functions of MicroRNAs in cancer, mismatch repair, folate biosynthesis, fatty acid metabolism, and peptidoglycan biosynthesis function were decreased in the train driver group relative to the official group (Fig. 5A). While in the prediction of fungal function, there were significant differences between the official group and the train driver group in Trans-2-enoyl-CoA reductase (NADPH), Beta-ureidopropionase, choline kinase, Catechol O-methyltransferase, Cysteine dioxygenase, Arylsulfatase, Manan endo-1-6-alpha-mannosidase, Tropinone reductase | and Alkaline phosphatase. (*P* ⩽ 0.05). All functions above were decreased in the train driver group compared with the official group (Fig. 5B).

**Fig. 5.**
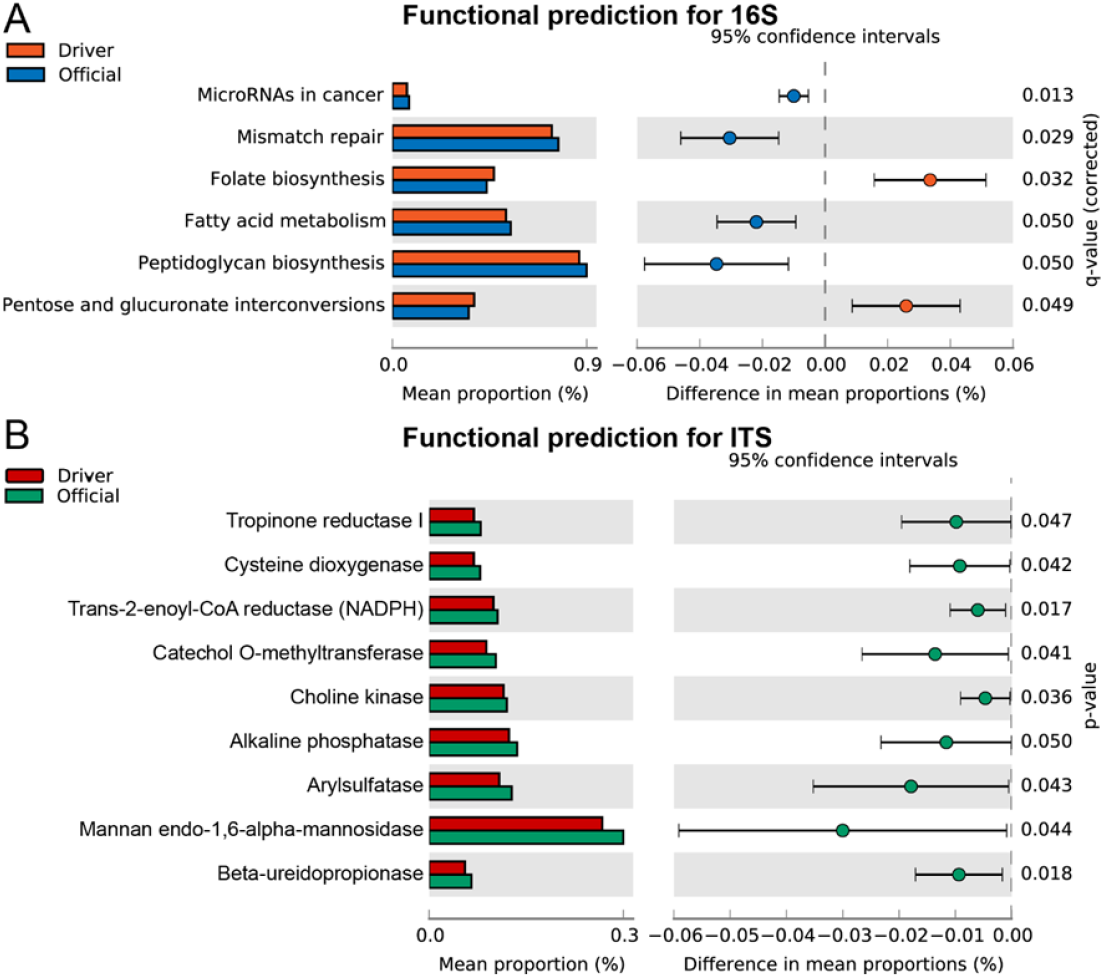
Functional predictions for the gut microbiota. A, The KEGG pathway with significantly different abundances in the two groups; B, The KEGG enzyme with significantly different abundances in the two groups. The left figure shows the abundance ratio of different functional classifications in the two groups. The middle figure shows the difference ratio of functional classifications abundance within the 95% confidence interval. The rightmost value is *P* value, and *P* value < 0.05 indicates significant difference.

## 4. Discussion

Our results indicates that the community richness and diversity of the official group were higher than those of the train driver group (Fig. 1A and 1B). Carlo Bressa *et al*. [10] found that a sedentary lifestyle significantly affects the type of microorganisms that promote health in women. Other studies [11] have proved that the elderly’s physical weakness and poor exercise ability are related to low fecal microbial diversity. In addition, the gut microbiota diversity of professional athletes is higher than that of people with the same BMI [12]. All the above studies have proved that both sedentary lifestyle and physical exercise can affect the gut microbiota diversity. Therefore, we speculated whether the difference in Alpha diversity of gut microbiota between the official group and the train driver group was due to the difference in sedentary lifestyle and physical exercise. As shown in Table 1, the sedentary rate of the train driver group was significantly higher than that of the offical group(P<0.001) and there was no significant difference in exercise between the two groups (P>0.5). Therefore, the differences in gut microbiota diversity between the two groups may be due to the differences in sedentary conditions.

It is widely accepted that a highly diverse and stable microbiome promotes overall human health. Decrease in intestinal microbiota Alpha diversity has been shown to be associated with a variety of acute and chronic diseases[13], such as inflammatory enteritis, helicobacter pylori infection, colon cancer, metabolic syndrome, type I and type II diabetes, cardiovascular disease, allergy, asthma, eczema, and autism. Multiple studies have shown that the fungal diversity of patients with cystic fibrosis (CF) is lower than that of healthy people[14]; Sonja Lang[15]et al. found that patients with alcohol-related liver disease had lower fungal diversity; Bajaj[16]et al. demonstrated that fungal dysbiosis and *Candida* overgrowth in 143 cirrhosis patients. Intestinal fungal dysbiosis has also been observed in other diseases with intestinal barrier dysfunction, such as IBD[17]. Although the pathogenic role of fungi in IBD has not been clearly demonstrated[18], fungal dysbiosis contributes to ethanol-induced steatohepatitis in mice. Therefore, maintaining the stability of intestinal microbiota diversity is important for human health.

NMDS (Nonmetric Multidimensional Scaling) analysis showed that microbial community compositions of the official group and the train driver group were similar in (Fig. 1C,D). Although there were significant difference between the two groups in the microbial diversity and richness, and specific microorganisms, the two groups are both from China railway corporation, who are living in similar environment and owning the same diet structure.There were no difference in exercise and diet (table 1), so two groups’ similarity in microbial community composition is reasonable.

Our research results showed that the dominant phylum of the two group were Fimicutes, Bacteroidota, Actinobateriota and Protecbacteria, among which Firmicutes accounted for more than 70% of the total (Figure 2C). Studies have shown that the gastrointestinal microbiota is mainly composed of Firmicutes, Bacteroidetes, Proteobacteria and Actinobacteria [5]. Therefore, our findings are consistent with previous studies. Sequencing analysis of fecal samples showed that there were great differences among healthy individuals: Bacteroidetes and Firmicutes showed great differences in relative abundance among 242 healthy individuals [19]. Other studies [20-21] found that the main axis of intestinal microbiome variation was Firmicutes -Bacteroidetes axis. In our study, the difference in phylum levels between the two groups was mainly in Bacteroidetes and Firmicutes, which was consistent with previous studies. On genus level, the gut microbiota of the official group and the train driver group was composed of *Faecalibacterium, Blautia, Bacteroides, Megamonas, Bifidobacterium, Agathobacter, Escherichia*-*Shigella, Subdoligranulum, Roseburia, Preverella Votella*), Unclassified *Lachnospiracee, Eubacterium Hali Group* and others. Both groups had similar community composition on phylum and genus levels, which further supported the results of NMDS analysis.A comparative analysis of fecal microbiota in Chinese isolated Yao population, minority Zhuang and rural Han by 16S RNA gene sequencing showed that on genus level, all three groups were mainly composed of *Bacteroides, Prevotella, Faecalibacterium, Roseburia, Lachnospira, Sutterella, Ruminococcus, Parabacteroides, Coprococcus* and *Oscillospira* [22]. In a recent study of healthy Italians who were followed for more than a year, the dominant phyla of intestinal microbiota were *Firmicutes* and *Bacteroidetes*, followed by *Proteobacteria, Actinobacteria*, and *Verrucomicrobia*. At the family level, the dominant bacteria were *Ruminococcaceae, Lachnospiraceae, Bacteroidaceae* and *Prevotellaceae*, while at the genus level, the dominant were *Bacteroides, Pervotella* and *Faecalibacterium* [23]. Our results are in part consistent with previous studies, and discrepancies may be due to differences in diet and geographical location.

The ITS sequencing analysis showed that at phylum level, the gut fungi in the official group and the train driver group were mainly composed of *Ascomycota* and *Basidiomycota*. On genus level, the two groups were both composed of *Candida, Aspergillus, Penicillium, Apiotrichum, Cladosporium, Cryptococcus, Monascus*, etc. The distribution of fungi showed significant differences in distant body parts, while similar distribution patterns were found in adjacent parts. Several studies using next-generation sequencing have identified different fungal communities in the human gut, mainly including *Ascomycota, Basidiomycota*, and *Zygomycota* [24-26]. One of the studies identified 16 species of fungi in fecal samples [27], with *Galactomyces geotrichum* being the most prevalent. Another study revealed 66 species of fungi, of which *Saccharomyces* was the most common, followed by *Candida* and *Cladospora* [24], while another study found 75 species of fungi, of which *Penicillium, Candida* and *Saccharomyces* were the most common [26]. The fecal analysis of 96 healthy individuals by Hoffmann et al. identified 66 genera of fungi, the most common being *Saccharomyces, Candida* and *Cladospora*. As mentioned above, a wide variety of fungi have been found in the human gastrointestinal tract, but there is no consensus on the ideal gut fungal microbiome.

There were significant differences in relative abundance of some taxa microbiota between the two groups. LEfse analysis showed that the relative abudance of *Roseburia* in the official group was higher than that of the train driver group. Bressa, C[10] et al. found more health-promoting bacteria species in active women, including *Faecalibacterium prausnitzii, Roseburia hominis* and *Akkermansia muciniphila*.The analysis of Castellanos N[28] et al. also suggests that *Roseburia* is a key centre for a active lifestyle. The presence of *Roseburia* is associated with more frequent intestinal transport, which may also explain why the higher abundances of *Roseburia* appear in active lifestyles, as people with active lifestyles generally have more active intestinal transport.So the higher relative abundance of *Roseburia* in the official group may be related to lower rate of sitting compared with the train driver group.

The receiver-operating characteristic (ROC) curves showed that these differential microbiota by LEfSe analysis can distinguish between the two groups with accuracy and effectiveness, indicating that there are significant difference in gut microbiota of various populations.The differences may be related to characteristics of different occupations.

In our study, we found that some bacteria and fungi were correlated with clinical characteristics, indicating that the gut microbiota are impacted by a variety of factors.Moreover, function prediction suggested that some genera were correlated with physiological fuctions.Considering that metagenomics can provide more detailed information, especially functional analysis and deeper analysis of species. Therefore, further metagenomic studies should be carried out according to the research results.

Interactions between the fungal and bacterial microbiome may also play a role in health and disease. In some cases, the presence of bacteria was positively correlated with the presence of fungi. For example, *Mycobacterium* reinfection sometimes occurs with *Aspergillus* [29]. In other cases, bacteria compete with fungi; When *Pseudomonas aeruginosa* was dominant in CF[30], the growth of *Candida* and other fungi was inhibited. Therefore, further studies can be made on the relationship between bacteria and fungi.

Previous studies mostly focused on the differences in fecal microbiota among people of different races or geographical locations. Our study firstly analyzed the differences in gut microbiota among people of different occupations in the same region, which is somewhat innovative. Questionnaire survey made a detailed understanding of the subjects’ basic information, dietary habits, living habits, family history and past medical history, etc. Therefore, many factors were considered in the analysis of intestinal microbiota. However, there are still some limitations in this study: a total of 172 railway workers were involved in this study, which were divided into 2 groups, and the population and occupation types could be expanded in the future. Most of the subjects in this study were male, so it was difficult to compare the effects of male and female genders on microbial composition and function. In addition, there is a lack of serological test results to study the relationship between biochemical indicators, immune status and intestinal microbiota.

## Conclusion

These results demonstrate that the gut microbiota between officials and train drivers were different in some aspects, indicating that the microbiota may be directly affected by a dynamic working environment. Further metagenomic studies should be carried out to get more functional analysis and deeper analysis of species.

## Acknowledgements

We thank the subjects who provided their fecal samples.This research was supported by China Railway. We declare that we have no competing interests.

## Supplementary materials

**Fig. S1.**
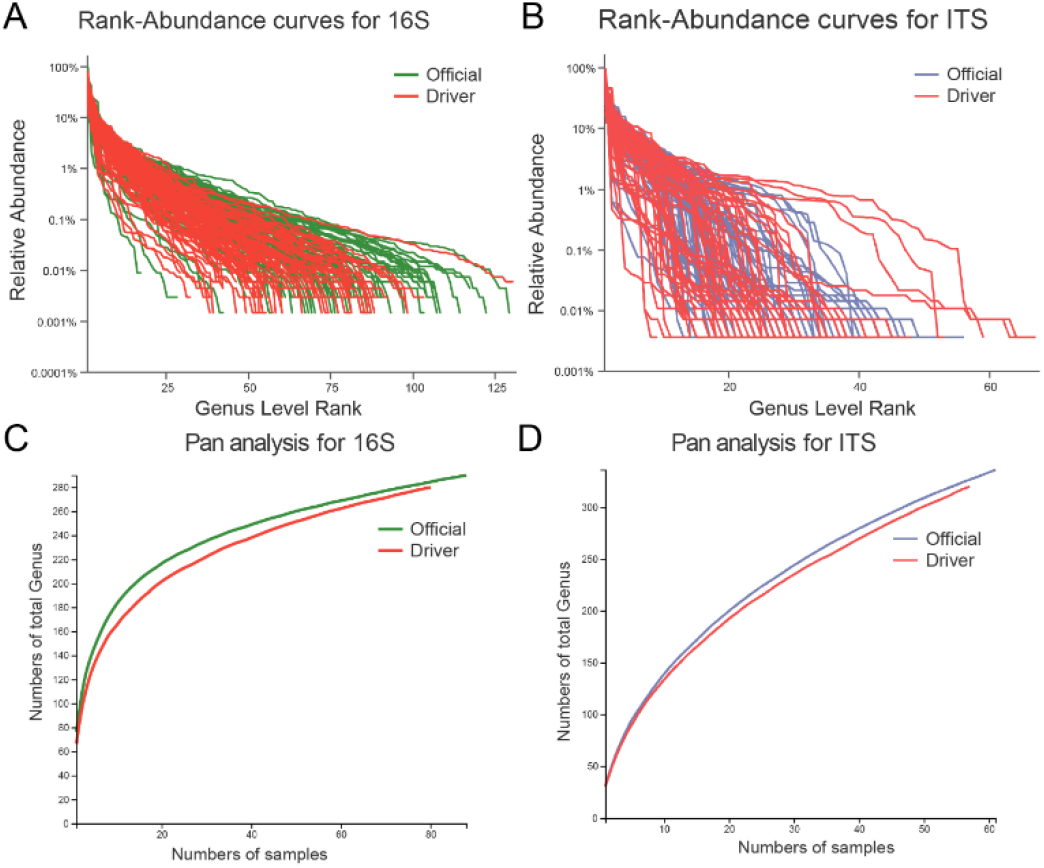
Relative abundance of the gut microbiota. A, Rank-abundance curves for the bacterial gut microbiota on the genus level; B, Rank-abundance curves for the fungal gut microbiota on the genus level; C, Number of genera for the bacterial gut microbiota; D, Number of genera for the fungal gut microbiota.

**Fig. S2.**
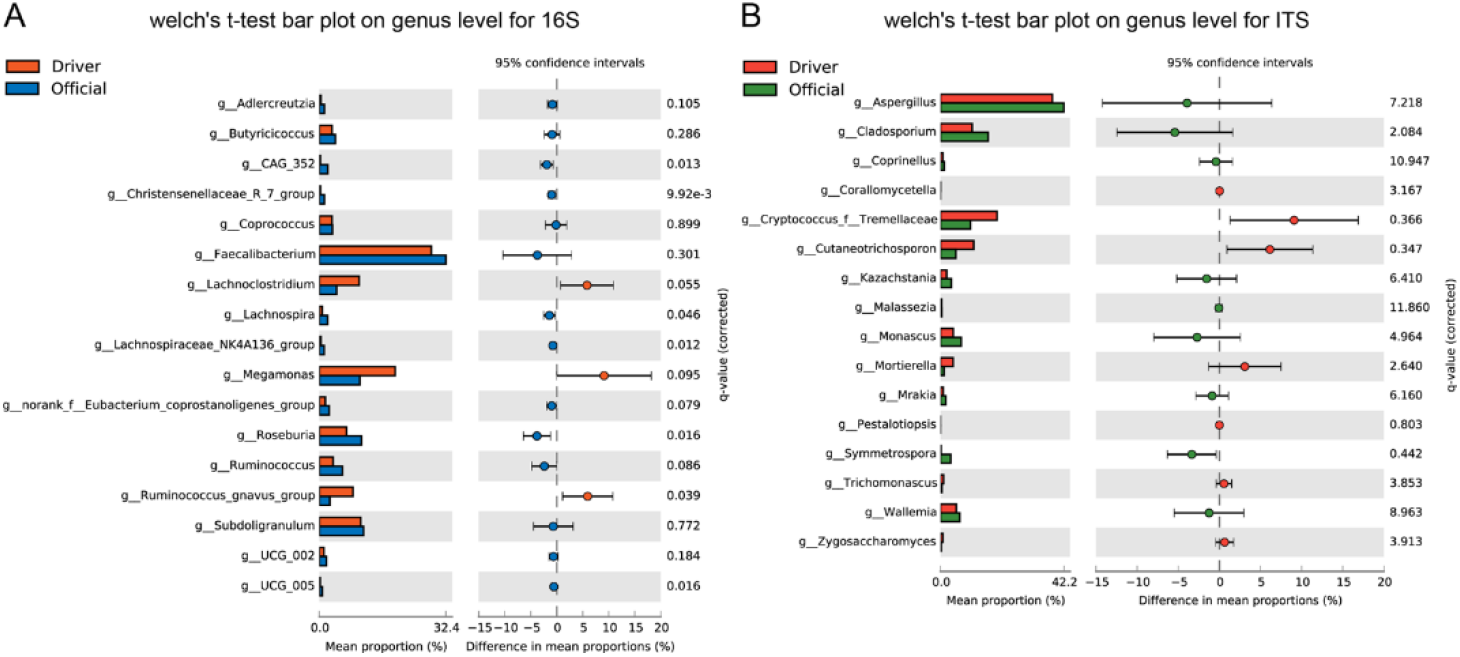
The abundance of different species in LEfSe analysis. A, The abundance of the bacterial gut microbiota for two group in genus level; B, The abundance of the fungal gut microbiota for two group in genus level.

## References

[1] Lagier JC, Dubourg G, Million M, et al. Culturing the human microbiota and culturomics. Nat Rev Microbiol, 2018,16:540–550.

[2] Dominguez-Bello MG, Costello EK, Contreras M, et al. Delivery mode shapes the acquisition and structure of the initial microbiota across multiple body habitats in newborns. Proc Natl Acad Sci U S A. 2010, 107(26):11971–11975.

[3] Turnbaugh PJ, Ley RE, Hamady M, Fraser-Liggett CM, Knight R, Gordon JI. The human microbiome project. Nature. 2007,449(7164):804–810.

[4] Sam QH, Chang MW, Chai LY. The Fungal Mycobiome and Its Interaction with Gut Bacteria in the Host. Int J Mol Sci. 2017 Feb 4;18(2):330.

[5] Quaranta G, Sanguinetti M, Masucci L. Fecal Microbiota Transplantation: A Potential Tool for Treatment of Human Female Reproductive Tract Diseases. Front Immunol. 2019, 10:2653.

[6] Huffnagle GB, Noverr MC. The emerging world of the fungal microbiome. Trends Microbiol, 2013,21(7):334–341.

[7] Francescangeli F, De Angelis ML, Zeuner A. Dietary Factors in the Control of Gut Homeostasis, Intestinal Stem Cells, and Colorectal Cancer. Nutrients, 2019, 11(12):2936.

[8] Cohen, R.; Roth, F.J.; Delgado, E.; Ahearn, D.G.; Kalser, M.H. Fungal flora of the normal human small and large intestine. N. Engl. J. Med. 1969, 280, 638–641.

[9] Becattini S, Taur Y, Pamer EG. Antibiotic-Induced Changes in the Intestinal Microbiota and Disease. Trends Mol Med, 2016,22(6):458–478.

[10] Bressa C, Bailén-Andrino M, Pérez-Santiago J, et al. Differences in gut microbiota profile between women with active lifestyle and sedentary women. PLoS One, 2017,12(2):p e0171352.

[11] Claesson MJ, Jeffery IB, Conde S, et al. Gut microbiota composition correlates with diet and health in the elderly. Nature, 2012, 488(7410):178–184.

[12] Clarke SF, Murphy EF, O’Sullivan O, et al. Exercise and associated dietary extremes impact on gut microbial diversity. Gut, 2014,63(12):1913–1920.

[13] Pickard JM, Zeng MY, Caruso R, et al. Gut microbiota: Role in pathogen colonization, immune responses, and inflammatory disease. Immunol Rev, 2017, 279(1):70–89.

[14] Nelson A, De Soyza A, Bourke S, Perry J, Cummings S: Assessment of sample handling practices on microbial activity in sputum samples from patients with cystic fibrosis. Lett Appl Microbiol, 2010, 51:272–277.

[15] Lang S, Duan Y, Liu J, et al. Intestinal Fungal Dysbiosis and Systemic Immune Response to Fungi in Patients With Alcoholic Hepatitis. Hepatology, 2020, 71(2):522–538.

[16] Bajaj JS, Liu EJ, Kheradman R, Fagan A, Heuman DM, White M, Gavis EA, et al. Fungal dysbiosis in cirrhosis. Gut, 2018,67:1146–1154.

[17] Sokol H, Leducq V, Aschard H, Pham HP, Jegou S, Landman C, Cohen D, et al. Fungal microbiota dysbiosis in IBD. Gut, 2017,66:1039–1048.

[18] Ni J, Wu GD, Albenberg L, Tomov VT. Gut microbiota and IBD: causation or correlation? Nat Rev Gastroenterol Hepatol, 2017,14:573–584.

[19] Cremer J, Arnoldini M, Hwa T. Effect of water flow and chemical environment on microbiota growth and composition in the human colon. Proc Natl Acad Sci U S A. 2017 Jun 20;114(25):6438–6443.

[20] Wu GD, Chen J, Hoffmann C, et al. Linking long-term dietary patterns with gut microbial enterotypes. Science, 2011,334(6052):105–108.

[21] Arumugam M, Raes J, Pelletier E, et al. Enterotypes of the human gut microbiome. Nature, 2011,473(7346):174–180.

[22] Liao M, Xie Y, Mao Y, et al. Comparative analyses of fecal microbiota in Chinese isolated Yao population, minority Zhuang and rural Han by 16sRNA sequencing. Sci Rep, 2018, 8(1):1142.

[23] Barone M, Turroni S, Rampelli S, et al. Gut microbiome response to a modern Paleolithic diet in a Western lifestyle context. PLoS One, 2019, 14(8):e0220619.

[24] Hoffmann, C.; Dollive, S.; Grunberg, S.; Chen, J.; Li, H.; Wu, G.D.; Lewis, J.D.; Bushman, F.D. Archaea and fungi of the human gut microbiome: Correlations with diet and bacterial residents. PLoS ONE, 2013, 8, e66019.

[25] Suhr, M.J.; Hallen-Adams, H.E. The human gut mycobiome: Pitfalls and potentials—A mycologist’s perspective. Mycologia 2015,107:1057–1073.

[26] MarRodríguez M, Pérez D, Javier Chaves F, et al. Obesity changes the human gut mycobiome. Sci Rep. 2015,5:14600.

[27] Hamad I, Sokhna C, Raoult D, Bittar F. Molecular detection of eukaryotes in a single human stool sample from Senegal. PLoS One. 2012,7(7):e40888.

[28] Castellanos N, Diez GG, Antúnez-Almagro C, et al. Key Bacteria in the Gut Microbiota Network for the Transition between Sedentary and Active Lifestyle. Microorganisms, 2020,8(5):785.

[29] Darling WM. Co-cultivation of mycobacteria and fungus. Lancet. 1976,2(7988):740.

[30] Kerr J. Inhibition of fungal growth by Pseudomonas aeruginosa and Pseudomonas cepacia isolated from patients with cystic fibrosis. J Infect. 1994 May;28(3):305–10.

